# Fast NeighborNet: Improving the Speed of the Neighbor-Net Phylogenetic Network Algorithm with Multithreading and a Relaxed Search Strategy

**DOI:** 10.1101/283424

**Authors:** Jacob Porter

**Affiliations:** Department of Computer Science, Virginia Tech, Blacksburg, VA 24061 USA

## Abstract

Fast Neighbor-Net is a command-line Java program that has fast implementations of the popular Neighbor-Net phylogenetic split network approach. This allows more efficiency in computationally intensive tasks such as larger scale data analysis and bootstrapping. The canonical search phase iteratively finds a pair of connected components that minimizes a distance function in Θ(*n*^3^) time in the input taxa count. A relaxed search strategy has been implemented that has average-case time complexity of *𝒪*(*n*^2^ log *n*) but with Θ(*n*^3^) worst-case time complexity. This search strategy sacrifices some accuracy for speed. The original approach’s implementation has been improved by using good programming practice. These improvements increased run-time performance by a constant factor of approximately 2 and reduced memory requirements by a constant factor of approximately 6. These search strategies allow multithreading to better use modern CPU hardware. PFAM data of 2000–30,000 taxa were used for testing performance. The canonical implementation with three threads improved average performance by approximately 2.1. The relaxed search has good quality, and the accuracy was tested on a mammal and a eukaryote data set. Kendall tau distance was used as a rough measure of topological similarity for the relaxed and canonical search strategies.

## 1. Introduction

The popular Neighbor-Net method of Bryant and Moulton is a phylogenetic split network algorithm with good speed and memory requirements [4] and proven statistical consistency [5]. The networks that it produces are planar with outer nodes labeled as taxa; thus, the networks are easy to draw and read. The network can represent recombination, hybridization, gene fusion, sequencing error, lateral gene transfer, and other signals of conflicting evolution that are not well represented by a tree [3]. These phenomena can be represented with a graph where some nodes have multiple parents unlike a tree where nodes have a single parent [11]. Neighbor-Net was inspired by the popular Neighbor-Joining algorithm [20].

Neighbor-Net takes as input a distance matrix and works by agglomerating connected components in an execution graph based on a distance function for each iteration. These terms will be defined in a later section. The original implementation is included in the SplitsTree4 Java program [10].

For *n* taxa, the Neighbor-Net algorithm has worst-case time complexity Θ(*n*^3^) and space complexity Θ(*n*^2^). Thus, it runs in seconds or minutes for input sizes of only a few thousand taxa but takes hours if not days for input sizes of tens of thousands or hundreds of thousands of taxa. Protein families of this size exist in the PFAM database [9]. Phylogenies of this size are useful in understanding questions in comparative genomics such as tree of life projects [18].

### Related Work

Neighbor-Joining and Neighbor-Net are very similar in their implementation details. There are several approaches for Neighbor-Joining [20] that have better average-case run-time. Methods that compute the same Neighbor-Joining tree faster than the canonical approach include QuickTree [13], QuickJoin [19], NINJA [24], and RapidNJ [23]. These methods have the same worst-case time complexity of Θ(*n*^3^) but require more memory because of their specialized data structures. NINJA, for example, uses an array of priority queues and increases memory requirements by a constant factor. Clearcut [22], a faster implementation of the Relaxed Neighbor-Joining approach [7], uses a relaxed search heuristic to achieve *𝒪*(*n*^2^ log *n*) average-case time complexity, but it is less accurate than the canonical approach. Another algorithm that computes phylogenetic split networks is split decomposition [1], but networks may not be planar and may be less resolved compared to Neighbor-Net [11]. Another popular split network is median networks [2].

## Contributions

To the author’s knowledge, there are no implementations of the Neighbor-Net approach that improve the run-time of the original implementation in SplitsTree in any way [10, 4]. Fast Neighbor-Net includes a fast search heuristic called relaxed search inspired by Relaxed Neighbor-Joining [7] and Clearcut [22], and it improves upon the run-time of the canonical approach. These implementations are faster in the following senses.

The run-time of the implementation of the canonical approach found in SplitsTree [10] was reduced by approximately half by using good programming practice. The memory requirements were reduced by about 9 for the quadratic part of memory. A relaxed search heuristic inspired by Clearcut for Neighbor-Joining was implemented. This gives *𝒪* (*n*^2^ log *n*) average-case time complexity but Θ(*n*^3^) worst-case time complexity. The relaxed search strategy reduces the accuracy of the algorithm. The canonical and relaxed search strategies employ multithreading. Computing the circular order faster makes analyzing larger data sizes possible and is useful for bootstrapping. The software “Noisy,” for example, computes circular orderings with Neighbor-Net to search for columns in a multiple sequence alignment that are phylogenetically less useful [6].

## 2. The Canonical Neighbor-Net Algorithm

Before discussing the Neighbor-Net algorithm in detail, several terms will be defined and a high level description of the algorithm will be provided.

### Definitions

The *taxa* set *𝒳* = {*x*_1_, …, *x*_*n*_} is a set of strings over an alphabet ∑, where each taxon *x*_*i*_ ∈ *𝒳* represents a group of organisms, such as a species, or an individual. For example, the string *x*_*i*_ could be the consensus sequence of a homologous protein, or each *x*_*i*_ could represent a different mammal such as cow, dog, gorilla, etc. In this example, the alphabet ∑ are symbols representing amino acids. The cardinality of *𝒳* is *n*.

A *split s* = {*A, B*} is a partition of the taxa *𝒳* into *A* and *B* so that *A* ∪ *B* = *𝒳* and *A*∩ *B* = ø. The *split weight function f_S_*: *S* → ℝ^+^ assigns a nonnegative real value to each split *s* ∈ *S*.

A metric *δ*_*X*_ on *𝒳* gives an estimate of the evolutionary distance between two taxa with respect to some evolutionary model. The metric space is (*𝒳, δ_𝒳_*). For biological sequence data, this metric can be computed with evolutionary models such as the Jukes-Cantor model [12], the Kimura parameter models [15], and other models [8]. The input to Neighbor-Net is the distance *δ*_*𝒳*_(*x, y*) for each distinct *x, y*∈ *𝒳*

During the execution of Neighbor-Net, the algorithm produces a *circular ordering, 𝒵* = {*z*_1_, *z*_2_,…, *z*_*n*_}, of the taxa. A *circular ordering* is a free cyclic permutation of the taxa.

### High Level Description

The Neighbor-Net algorithm has the following input and output:

### Input

A lower triangular matrix 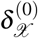 of pairwise taxon distances.

### Output

A circular ordering *𝒵* of the taxa, and a set of splits *S* together with a split weight function *f*_*S*_.

The algorithm consists of two phases: a search and agglomeration phase and a split weight estimation phase. The result of the search and agglomeration phase is a *circular ordering 𝒵* of the taxa. The ordering *𝒵* = *{z*_1_, *z*_2_, …, *z*_*n*_} where each *z*_*i*_ ∈ *X* uniquely defines a *split set S* given by

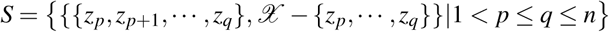

The splits together with their weights are the output of the split weight estimation phase of Neighbor-Net. This phase terminates the Neighbor-Net algorithm.

### Data Structures

During the execution of the search and agglomeration phase of Neighbor-Net, the algorithm maintains a set of nodes and two sets of edges on these nodes. This induces two graphs 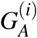, an agglomeration history graph consisting of a directed acyclic graph, and 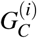, an undirected graph of connected components. As the algorithm proceeds, connected components from 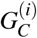 are combined in an operation called agglomeration, and the history of these operations is maintained in 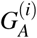 and an agglomeration history stack 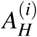. The purpose of this graph and the stack is to construct the circular ordering of the taxa. The two graphs are initially identical and consist of one node for each taxon input into Neighbor-Net and no edges. The agglomeration history stack is initially empty.

For each iteration, the graph 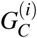 consists of two kinds of connected components: couples and singles. Couples are two connected nodes, and singles are singleton unconnected nodes. Because of the simplicity of 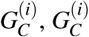 is stored as an array of node objects. The graph 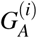 consists of node objects with references to sibling and children node objects.

The algorithm maintains an integer *m*^(*i*)^ that gives the number of connected components in 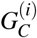. As the algorithm proceeds, the integer *m*^(*i*)^ gets smaller as connected components are combined. The Neighbor-Net algorithm maintains the distance matrix 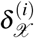 that gives the distances between nodes in 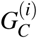. It is initialized to the distances given as input to the algorithm. The distance 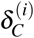 gives the distances between connected components in 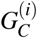 and is computed as needed according to a formula given in the next section. Row sums for 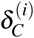 are maintained and stored as a field in a node representing a connected component.

An *agglomeration event* is an operation where two connected components are combined in 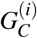 and replaced by a new connected component. The algorithm chooses two connected components that have the least distance to each other according to a function that will be explained in the next section. The agglomeration history graph, 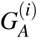, maintains a history of agglomeration events performed on the connected components graph. Because connected components consist of either a single or a couple, there are three possible agglomeration events: single-to-single, single-to-couple, and couple-to-couple. These are shown in Figure 1, and each case will be explained.

**Figure 1:**
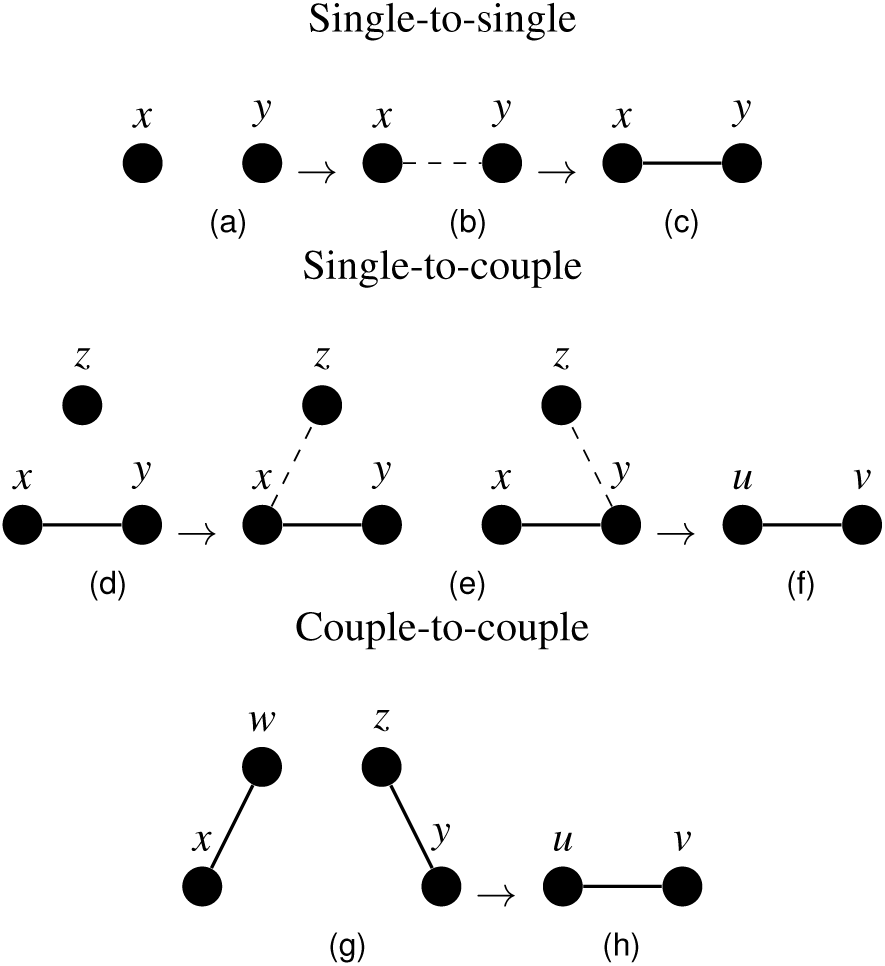
The three different agglomeration events performed in the agglomeration phase of Neighbor-Net. (a) Two singles are merged. (b) There is only one possibility, so (c) a couple is produced. (d) A single and a couple are combined, and (e) there are two possibilities to consider (f) resulting in a couple with new nodes. (g) A couple and a couple are combined with four possibilities (not shown) to examine (h) that results in a couple with new nodes.

For a single-to-single agglomeration, the algorithm defines a new edge between *x* and *y* in 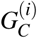, and a new directed edge from *x*, the lexicographically least node, to *y* in 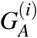. For a single-to-couple agglomeration, there is a single node *x* and two connected nodes *y* and *z*. In 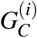, two new connected nodes *u* and *v* are created and *x, y*, and *z* are deleted. The node in the couple that is closest to *x* is found by a procedure described in the next section, and an edge from *x* to this node is created in the agglomeration history graph,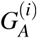. In the agglomeration history graph, the two new nodes *u* an *v* are connected with an edge going from *u* to *v*. An edge is connected from node *u* to *x* and from *u* to *x* the node closest to *x*. An edge is connected from node *v* to the node closest to *x* and to the node furthest from *x*. An edge goes from *x* to the node closest to *x*. The node *u*, the lexicographically least node for the two new nodes created, is added to the agglomeration history stack, *A*_*H*_.

In the last case, a couple consisting of nodes *w* and *x* is agglomerated with another couple consisting of *z* and *y*. Without loss of generality, suppose that the two nodes that are the closest to each other are *x* and *y*, then two single-to-couple agglomeration events are performed. The first agglomeration event is done on *w, x*, and *y* to produce *s* and *t* and then another agglomeration event is done on *s, t*, and *z* to produce *u* and *v*.

After an agglomeration event, the distances between the nodes 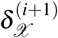 and the row sums are updated as described next.

### Computing Distances

The distance between connected components *c*_*j*_ and *c*_*k*_ in the connected components graph 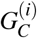 is the average distance between their constituent nodes. Thus, the connected component distance is given by

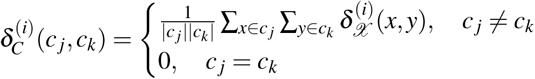

For a connected component *c*_*k*_ in 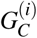, the row sum 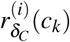 is the sum of connected component distances between *c*_*k*_ and all the other connected components.

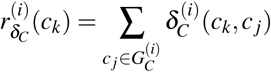

Let *m*^(*i*)^ be the number of connected components in the graph 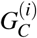. The connected component pair *c*_*k*_ and *c _j_* that minimizes the following function, called the *Q*-criterion, is chosen for agglomeration.

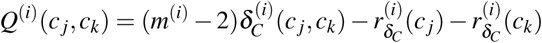

The *Q*-criterion has this form so that it is linear, permutation equivariant, and statistically consistent [5]. In the canonical implementation, it is calculated with a doubly nested `for` loop over all existing connected components. This gives a quadratic time complexity for each iteration. It is this part that is the focus of some of this work’s enhancements.

Now that two connected components *c*_*j*_ and *c*_*k*_ have been chosen, they must be agglomerated. If one of the connected components is a couple, the agglomeration history graph 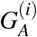 connects nodes with respect to nodes between connected components that have the least distance according to the 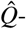criterion that will be described now.

The value 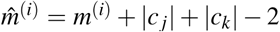 counts all the connected components in 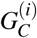 minus two plus all the nodes in components *c*_*j*_ and *c*_*k*_ This is as if all the nodes in are treated as singleton components. Treating nodes in *c*_*k*_ and nodes in *c*_*j*_ as if they were singleton nodes in the connected components graph, we wish to find the node *n*_*r*_ ∈ *c*_*k*_ and the node *n*_*s*_ ∈ *c*_*j*_ that minimizes the following

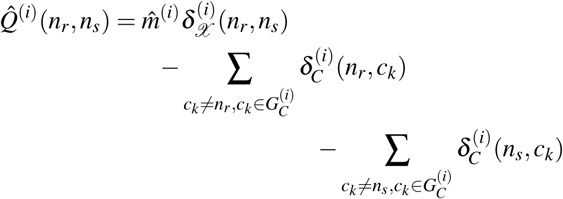

When agglomeration occurs, the distance matrix 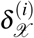 must be updated and reduced to represent fewer nodes and new distances between nodes. The updated distance, 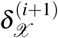, is computed in the following manner. If a single-to-single agglomeration event occurred,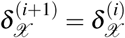. Suppose a single-to-couple agglomeration event occurred where *x* and *y* were a couple and *y* was the node with the least distance from the single node *z*. These nodes are replaced by nodes *u* and *v* where the distance from these nodes to a node 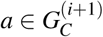 is given by the following formulas.

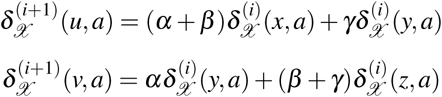

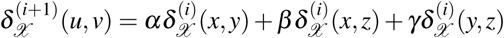

The parameters *α, β, γ* are constants such that *α* + *β* + *γ* = 1. In the canonical implementation,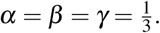.

### Circular Ordering Computation and Split Weight Estimation

The Neighbor-Net algorithm agglomerates connected components until there are only three nodes (two connected components) in the graph 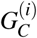. When this occurs, the algorithm constructs a simple undirected cycle 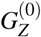 of the three nodes. The algorithm creates a circular ordering *𝒵* from this, the agglomeration history graph, and the agglomeration stack.

From the circular ordering *𝒵*, the set of splits *S* that respect the circular ordering can be calculated, and their weights can be calculated by using the least squares method. Ordinary least squares can produce negative split weights, which have no evolutionary meaning, so a non-negative constraint is imposed. The circular split network is mathematically motivated and allows the phylogenetic network to be visualized in a plane; however, some biological phenomena may not be able to be visualized in a plane, so the Neighbor-Net approach could miss those phenomena [4, 17]. Further details can be found in [11] and [4].

However, the constrained optimization problem requires substantial computational time. For an input of 2000 taxa, the circular ordering computation took several minutes while the split weight computation took three days. Currently, the split weights are estimated with a method by Lawson and Hanson [16] implemented in Parallel Java 2 [14]. However, this code takes *θ* (*n*^4^) memory. A customized solver that takes *θ* (*n*^2^) memory is being developed.

### Neighbor-Net Algorithm Summary

The algorithm 1 gives a summary of the neighbor net algorithm. The distance matrix *δ*_*X*_ is the input matrix, and the graphs 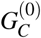 and 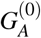 are initialized to a set of *m*^(0)^ = *n* unconnected nodes. The algorithm then iteratively agglomerates minimizing connected components in 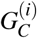 according to the *Q*-criterion storing the agglomeration history in 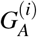 and the stack 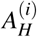. When there are only three nodes left, the `while` loop on line 6 terminates and the three remaining nodes are put into a simple cycle 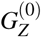. This graph is updated by popping off nodes from 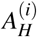 and using 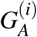 to find the neighbor and children of the nodes. The children in 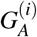 replace the parents in 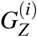 until the stack is empty. At this point, the order that the nodes are traversed in the simple cycle in 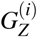 gives the circular ordering *𝒵* that determines the splits *S* on lines 14 and 15. Finally, in the split weight estimation phase on line 16, the split weights are calculated. The algorithm returns the circular ordering *𝒵*, the splits *S*, and the split weights *f*_*S*_. Visualizing the network can be done with the algorithm from section 7.2 of the book Phylogenetic Networks by Huson, Rupp, and Scornavacca [11]. Fast Neighbor-Net outputs a Nexus file that can be visualized in SplitsTree4.

#### Algorithm 1 Neighbor-Net

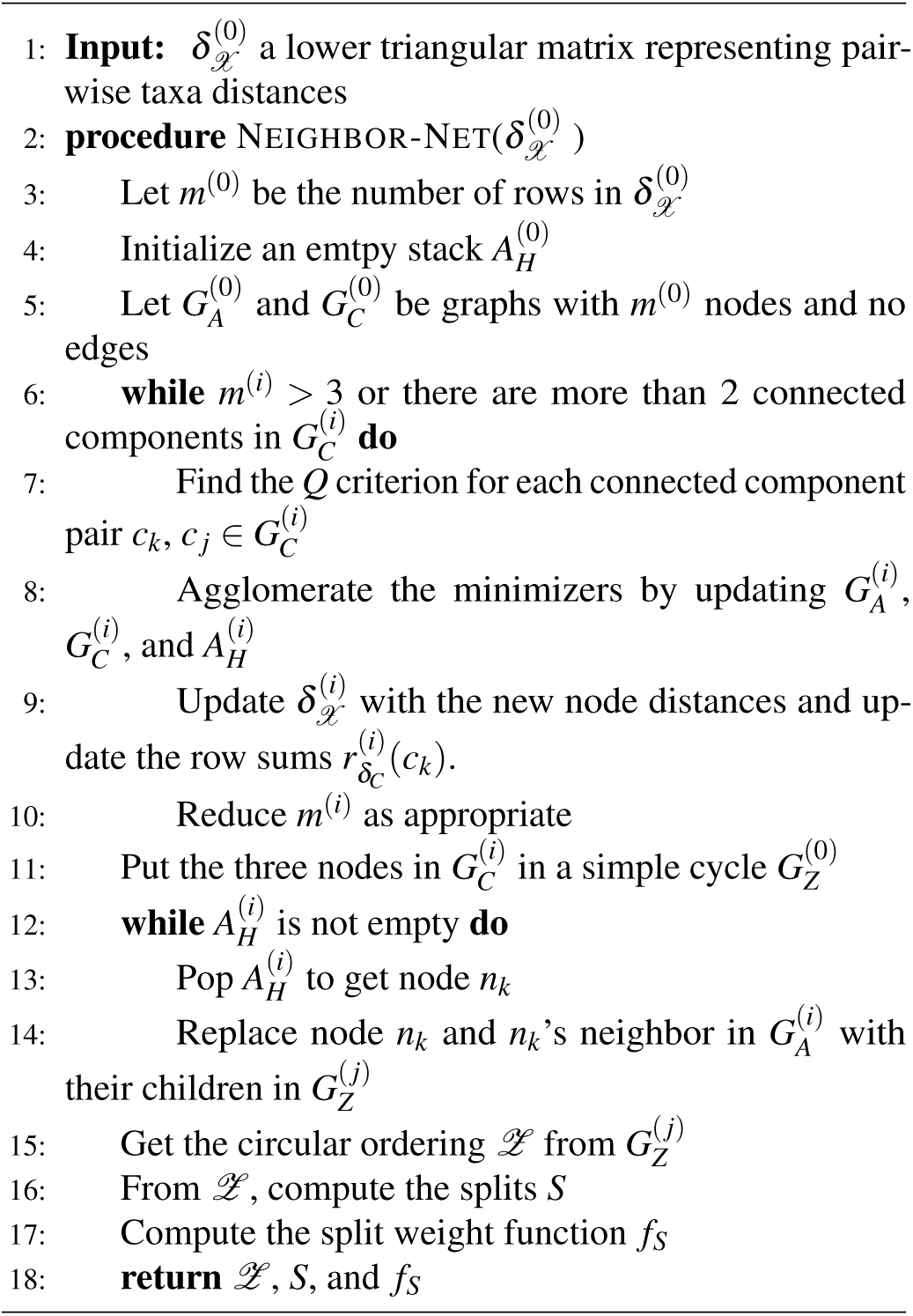

The time complexity of the search and agglomeration phase is Θ(*n*^3^). It can be shown that there are *θ* (*n*) iterations of the `while` loop on line 6, and there are *𝒪* (*n*^2^) pairs to calculate the *Q*-criterion on line 7. Lines 8 and 10 take constant time, and line 9, updating the distance matrix and row sums, takes linear time. Storing the distances takes *θ* (*n*^2^) memory.

## 3. The Relaxed Search Strategy

The canonical search strategy chooses a pair of connected components among all connected components in 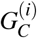 that minimizes the *Q*-criterion in line 7 of Algorithm 1. The relaxed search strategy replaces this approach by choosing a pair that minimize the *Q*-criterion with respect to each other.

More formally,

1. Until an agglomeration event occurs, select the next connected component *c*_*m*_ from a permutation 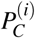 of the list of connected components in 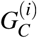. Do the following.
  a. Calculate *c*_*k*_ = argmin 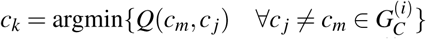
  b. Calculate *c*_*l*_ = argmin 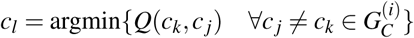
  c. If *c*_*m*_ = *c*_*l*_ agglomerate *c*_*m*_ and *c*_*k*_.

This heuristic has worst-case time-complexity Θ(*n*^3^) since all connected component pairs may need to be examined. In the best-case, step 1(a) will have an immediate minimizing neighbor and will not need to do more than one iteration of the `for` loop of line 1. According to the Relaxed Neigbhor-Joining paper [7], this occurs with probability ranging from 4*/n* to 1. The probability is higher for perfectly balanced trees and lower for pectinate “comb-like” trees. This gives an average-case time complexity of *𝒪* (*n*^2^ log *n*). The heuristic is nondeterministic, and calculating the permutation uses the unbiased Fisher-Yates procedure.

## 4. Results and Discussion

The correctness and accuracy of the canonical approach need not be tested since it has been tested in the existing literature [3, 4, 5], but the speed enhancements of this approach have been studied. The quality of the relaxed search strategy will be discussed.

### 4.1 Speed

Performance tests were performed on an Intel Core i7-5820k 6-core CPU @ 3.30 GhZ with 16 GB RAM and a 12GB Java heap in server mode on OpenSuse 13.2 and Java 8. Arbitrary protein family taxa were taken from the PFAM database for performance testing. There were around 50 files with between 2000–10000 taxa. The sequence data was converted to Phylip distance format using QuickJoin [19] and the process in [24].

The original Neighbor-Net code from SplitsTree4 was modified and integrated into Fast Neighbor-Net. This allows a fair comparison of speed. Fast Neighbor-Net calculates the row sums on line 9 in Algorithm 1 in linear time by subtracting the old distances from each connected component and then adding the distance to the new connected component to each connected component. The original implementation recreated the row sums from scratch, a quadratic process. Therefore, the single-threaded run-time of Fast Neighbor-Net is approximately twice that of the original. The distance matrix, 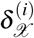, in Neighbor-Net used a row for each new node, but Fast Neighbor-Net reuses rows when new nodes are created. This reduced memory needed for the array by approximately 9.

Fast Neighbor-Net implements parallelism by giving each thread an equal-sized block of connected component pairs to compute the *Q*-criterion. Figure 2 shows how run-time scaled with 1, 3, and 5 threads. (Executions with 2 and 4 threads were excluded for readability.) Using three threads improved performance by an average of approximately 2.1. The relaxed approach was implemented in parallel by giving each thread a block of taxa pairs to compute the *Q*-criterion. The relaxed approach only showed performance gains with 10,000+ taxa.

**Figure 2:**
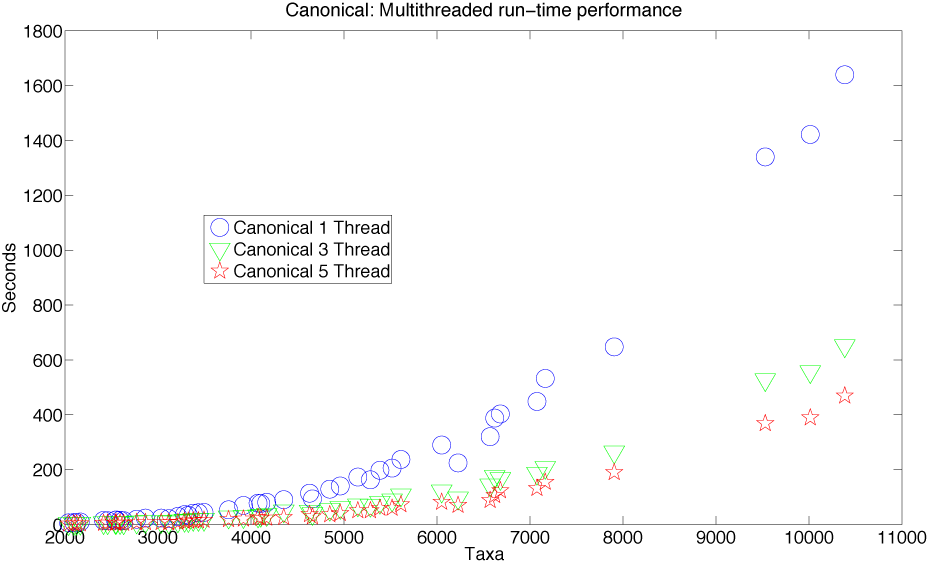
Multithreaded performance of the canonical approach.

### 4.2 Quality

#### Mammals and Branching Eukaryotes Data

Two real data sets were used to test the quality of the relaxed search approach. A 30 taxa set for mammals was taken from the SplitsTree4 software. Figure 3 shows that the major mammal clades are nicely resolved.

**Figure 3:**
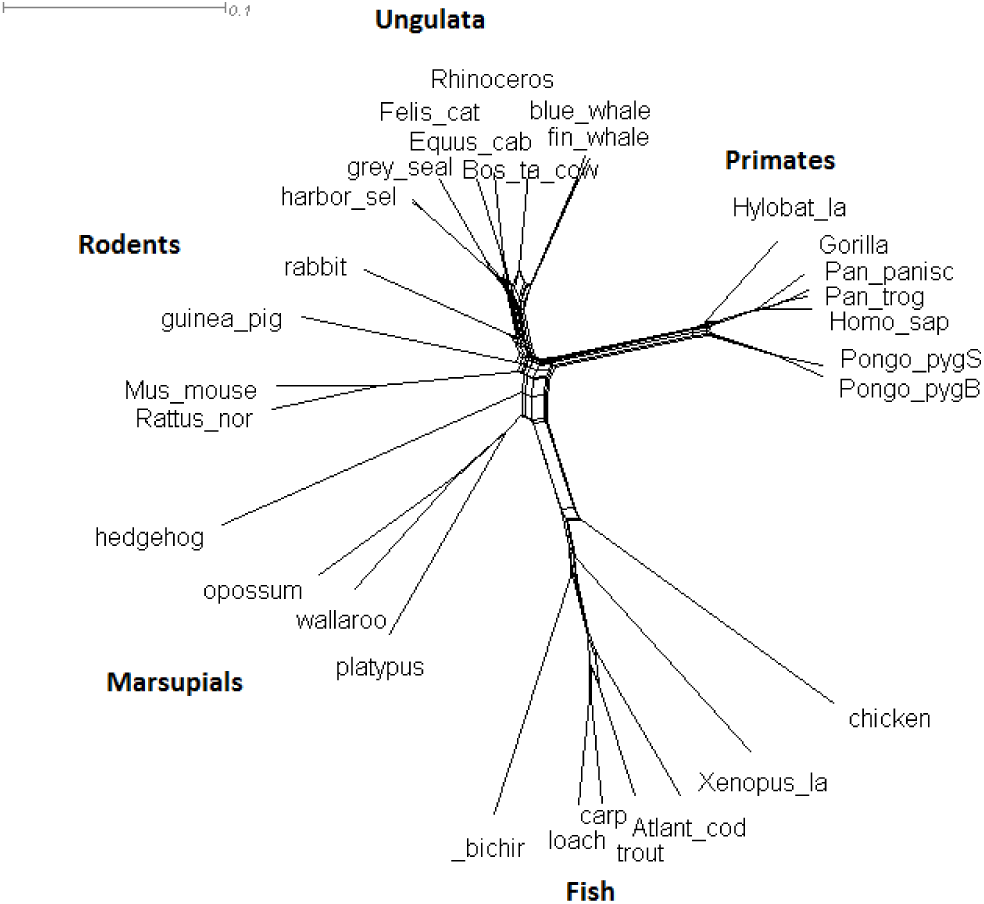
A thirty mammal phylogeny from data from SplitsTree4 computed with the relaxed search strategy. All the major clades are resolved accurately.

The evolution of early eukaryotes is an important part of evolutionary theory. Analyzing data with mitochondrial genes is challenging because of the few genes on the mitochondria, but it has led to a better understanding of evolutionary signal. The approach of [21] analyzed gene order data of mitochondria for 18 taxa using the normalized breakpoint distance. Biologists often use uncorrected distances like this when the evolutionary model is uncertain. Figure 4 shows a nice resolution of major clades. It resolved the green algae and red algae, and it grouped the alveolates with the stramenopiles. This is consistent with recent hypotheses. Neighbor-Joining and split decomposition did not do this [3]. The local search strategy produced the same result as the canonical strategy.

### Topology Distance

The circular ordering gives a general topology for the split network, so comparing the distance between the circular ordering computed by the canonical approach and the circular ordering compute by the relaxed heuristic gives an idea of whether the relaxed heuristic is better than a random topology. The normalized Kendall tau distance between the canonical approach’s ordering and the relaxed approach’s ordering was computed on 37 PFAM data sets of 2000–13000 taxa and averaged across five executions of the relaxed heuristic. The average was 0.379, and all distances were below 0.5, which is the average distance to a random permutation. This suggests that the topologies inferred are reasonably good. The least distance was 0.10, which is quite close.

**Figure 4:**
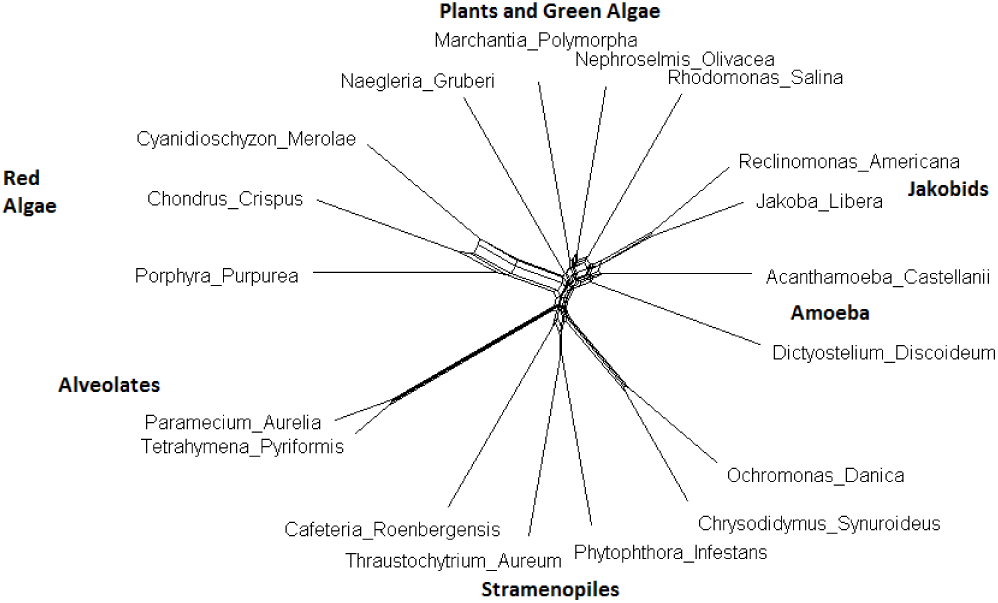
Early branching eukaryotes.

### 4.3 Correctness

Input distances are *additive* when they correspond to the unique distances between taxa defined by the edges of a tree. The canonical search strategy recovers this tree when the input distances are additive [4]. Relaxed Neighbor-Joining recovers the correct tree when the distances are additive and an additive check is performed [7]. Therefore, the relaxed search heuristic for Neighbor-Net should recover the correct tree when the input distances are additive and an additive check is performed.

## 5. Conclusion and Future Work

Fast Neighbor-Net is a fast implementation of the Neighbor-Net algorithm with a new relaxed search strategy that has good accuracy and performance qualities. Its input is a Phylip distance file, and it produces a Nexus file that can be rendered in SplitsTree4. It has a command-line interface for deployment on institutional compute clusters. The algorithm was modernized with multithreading. It is available at jacobporter.com/software/. Other search strategies such as a less accurate but faster random search strategy has been implemented. The memory requirements of the split weight estimation can be optimized. Further analysis is planned.

## Acknowledgements

David Bryant provided the original Java implementation of Neighbor-Net, and he provided the eukaryote data. Travis Wheeler provided the PFAM data. Layne Watson of Virginia Tech gave helpful comments about terminology and mathematical notation.

## References

[1] H.-J. Bandelt and A. W. Dress, “Split decomposition: A new and useful approach to phylogenetic analysis of distance data,” Molecular phylogenetics and evolution, vol. 1, no. 3, pp. 242–252, 1992.

[2] H.-J. Bandelt, V. Macaulay, and M. Richards, “Median networks: Speedy construction and greedy reduction, one simulation, and two case studies from human mtdna,” Molecular phylogenetics and evolution, vol. 16, no. 1, pp. 8–28, 2000.

[3] D. Bryant and V. Moulton, “Neighbornet: An agglomerative method for the construction of planar phylogenetic networks,” in Proceedings of the 2nd International Workshop on Algorithms in Bioinformatics. Springer, 2002, pp. 375–391.

[4] D. Bryant and V. Moulton, “Neighbor-net: An agglomerative method for the construction of phylogenetic networks,” Mol Biol Evol., vol. 21, pp. 255–65, 2004.

[5] D. Bryant, V. Moulton, and A. Spillner, “Consistency of the neighbornet algorithm,” Algorithms for Molecular Biology, vol. 2, pp. 8–20, 2007.

[6] A. W. Dress et al., “Noisy: Identification of problematic columns in multiple sequence alignments,” Algorithms Mol Biol, vol. 3, no. 7, pp. 7188–3, 2008.

[7] J. Evans, L. Sheneman, and J. Foster, “Relaxed neighbor joining: A fast distance-based phylogenetic tree construction method,” Journal of molecular evolution, vol. 62, no. 6, pp. 785–792, 2006.

[8] J. Felsenstein, Inferring Phylogenies. Sunderland, Mass.: Sinauer Associates, 2004.

[9] R. Finn et al., “The pfam protein families database,” Nucleic Acids Research, vol. 38, pp. D211–222, 2010.

[10] D. Huson and D. Bryant, “Application of phylogenetic networks in evolutionary studies,” Molecular Biology and Evolution, vol. 23, no. 2, pp. 254–267, 2006.

[11] D. H. Huson, R. Rupp, and C. Scornavacca, Phylogenetic Networks. Cambridge, United Kingdom: Cambridge University Press, 2010.

[12] T. H. Jukes and C. R. Cantor, “Evolution of protein molecules,” Mammalian protein metabolism, vol. 3, pp. 21–132, 1969.

[13] A. B. K. Howe and R. Durbin, “Quicktree: Building huge neighbour-joining trees of protein sequences.” Bioinformatics, vol. 11, pp. 1546–7, 2002.

[14] A. Kaminsky, “Parallel java: A unified api for shared memory and cluster parallel programming in 100% java,” in Parallel and Distributed Processing Symposium, 2007. IPDPS 2007. IEEE International. IEEE, 2007, pp. 1–8.

[15] M. Kimura, “A simple method for estimating evolutionary rates of base substitutions through comparative studies of nucleotide sequences,” Journal of molecular evolution, vol. 16, no. 2, pp. 111–120, 1980.

[16] C. L. Lawson and R. J. Hanson, Solving least squares problems. SIAM, 1974, vol. 161.

[17] D. Levy and L. Pachter, “The neighbor-net algorithm,” Advances in Applied Mathematics, vol. 47, no. 2, pp. 240–258, 2011.

[18] D. R. Maddison, K.-S. Schulz, and W. P. maddison, “The tree of life web project,” Zootaxa, vol. 1668, no. Linnaeus Tercentenary: Progress in Invertebrate Taxonomy, pp. 19–40, 2007.

[19] T. Mailund and C. Pedersen, “Quickjoin - fast neighbor-joining tree reconstruction,” Bioinformatics, vol. 20, pp. 3261–2, 2004.

[20] N. Saitou and M. Nei, “The neighbor-joining method: A new method for reconstructing phylogenetic trees.” Mol Biol Evol., vol. 4, pp. 406–25, 1987.

[21] D. Sankoff et al., “Early eukaryote evolution based on mitochondrial gene order breakpoints,” Journal of Computational Biology, vol. 7, no. 3-4, pp. 521–535, 2000.

[22] L. Sheneman, J. Evans, and J. Foster, “Clearcut: A fast implementation of relaxed neighbor joining,” Bioinformatics, vol. 22, pp. 2823–4, 2006.

[23] M. Simonsen, T. Mailund, and C. Petersen, “Rapid neighbor-joining,” in Proceedings of the 8th International Workshop on Algorithms in Bioinformatics. WABI, 2008, pp. 113–122.

[24] T. Wheeler, “Large-scale neighbor-joining with ninja,” in Proceedings of the 9th Workshop on Algorithms in Bioinformatics. WABI, 2009, pp. 375–389.

